# Assessing White Matter Engagement in Brain Networks through Functional and Structural Connectivity Mapping

**DOI:** 10.1101/2024.01.04.574259

**Authors:** Muwei Li, Kurt G Schilling, Zhaohua Ding, John C Gore

## Abstract

Understanding the intricate interplay between gray matter (GM) and white matter (WM) is crucial for deciphering the complex activities of the brain. While diffusion tensor imaging (DTI) has advanced the mapping of these structural pathways, the relationship between structural connectivity (SC) and functional connectivity (FC) remains inadequately understood. This study addresses the need for a more integrative approach by mapping the importance of the inter-GM functional link to its structural counterparts in WM. This mapping yields a spatial distribution of engagement that is not only highly reproducible but also aligns with direct structural, functional, and bioenergetic measures within WM, illustrating a notable interdependence between the function of GM and the characteristics of WM. Additionally, our research has uncovered a set of unique engagement modes through a clustering analysis of window-wise engagement maps, highlighting the dyanmic nature of the engagement. The engagement along with their temporal variations revealed significant differences across genders and age groups. These findings suggest the potential of WM engagement as a biomarker for neurological and cognitive conditions, offering a more nuanced understanding of individualized brain activity and connectivity patterns.

## Introduction

Brain function relies on the operations of distributed neural activities within the gray matter (GM), which do not operate in isolation but are intricately inter-connected to achieve complex functions. These interactions are facilitated by white matter (WM), which contains myelinated axonal fibers that create communication pathways between GM regions. The macro-architecture of these neural pathways has been extensively investigated, and with the development of diffusion MRI (dMRI), they may be reconstructed in vivo. Diffusion MRI also allows for the quantification of structural connectivity (SC) between two GM regions using diffusion metrics ^1–4^, such as the number of fibers that connect them, providing information on the strength and integrity of the neural conduits that underpin information exchange between GM areas. Concurrently, functional connectivity (FC) is assessed by analyzing the synchronization of activity across GM regions ^5,6^. This is often measured using functional MRI (fMRI) to detect blood oxygenation level-dependent (BOLD) signal fluctuations and their inter-regional temporal correlations. Previous imaging studies have suggested that although FC reflects SC to some extent, the relationship between them is complex ^7,8^, with some studies showing strong FC between regions that lack direct structural links, so that SC and FC may be complementary features of brain networks rather than being directly dependent ^9^. As a result, the integration of SC with FC has been the focus of numerous studies aiming to characterize brain organization more comprehensively. Some have reported that combining these features provides greater predictive accuracy regarding behavior or disease states than when each is used separately ^10–12^. Recent research has advanced beyond merely quantifying the level of SC to investigate its spatial topologies, particularly how WM extends into GM parcellations. This approach is particularly useful for evaluating the impact of WM lesions on GM activity, using known functional attributes of GM to predict the possible consequences of WM lesions on distinct brain functions ^13,14^.

While understanding the impact of extensions of WM structures into GM is of current interest, there has been little attention paid to assessing potential inverse effects by mapping functional measurements from GM onto WM. Such mappings could identify WM pathways critical for specific tasks or associated with functional deficits, especially when structural WM alterations are not apparent. Nozais et al. have pioneered the ‘functionnectome’, a methodology for integrating structural and functional data ^15^. This assigns values to WM voxels based on the weighted average of BOLD signals from all GM regions to which the voxel is structurally connected, with the weights being measures of SC.Thanks to the nature of the signals mapped from GM, the resulting data enable the application of standard fMRI analyses to WM. For example, using a general linear model (GLM), their studies have detected WM regions that are potentially involved in GM activation in response to tasks ^15^. Moreover, leveraging the segregated nature of the mapped signals, they have categorized WM voxels into distinct networks through independent component analysis (ICA) ^16^. However, the structural or biophysical foundations of these mapped BOLD signals remain unclear. In addition, given the growing evidence that BOLD signals within WM are measurable using appropriate analytical methods ^17–25^, the functional dependence between GM and WM can be more clearly illuminated by examining the relationships between mapped and directly measured BOLD signals in WM.

The ‘functionnectome’ model posits that changes in GM BOLD signals can be projected onto corresponding WM regions, assigning atypical BOLD responses to specific conditions such as disease or external factors. This model is especially powerful when investigating task-related BOLD changes. However, its application to resting-state data warrants caution due to the potential for bias. Averaging BOLD signals from various functional units during rest could introduce inaccuracies, as these baseline signals may not be as temporally synchronized as those elicited by tasks. This risk of bias is accentuated in WM areas containing crossing fibers, which appear as junctions for multiple, divergent functional systems. To address this, the present study builds on the same concept but maps specific graph metrics of the functional network instead of signals themselves to WM voxels. The process begins by modeling the brain as a graph wherein GM regions are considered as nodes, and edges represent the functional connections between these nodes. The edges are then analyzed for edge betweenness centrality (EBC), a metric indicating the importance of an edge as a bridge in the network. High betweenness centrality scores signify critical connectors whose removal could impede communication across the network. For each WM voxel, we trace the WM fibers with which the voxel intersects by using dMRI, and identify pairs of GM nodes that are linked by those fibers.

The value assigned to the WM voxel, named engagement, is the weighted average of the EBCs of the functional connections between the node pairs, where the weights are proportional to the SC. There are several benefits to this approach. Firstly, it circumvents the bias that may arise from averaging BOLD signals. Secondly, it maps the properties of functional edges directly to their structural counterparts (edge to edge), reflecting the same biological substrate rather than projecting nodal properties onto edges (node to edge). Thirdly, by using edge betweenness as a metric, we can assess how engaged each WM voxel is within the entire network, providing a more complete view of the role of WM voxels in brain connectivity. Our method offers a refined perspective by integrating functional and structural data, allowing for a more detailed description of the connectivity profile of the brain. This integration may be viewed as a supplement to our previous work, which examined WM engagement from a purely functional standpoint ^26^.

Our findings emphasize that the spatial distribution of WM engagement is highly reproducible, and correlates to a degree with myelination patterns, fiber structures, BOLD activity levels, and regional homogeneity, but also shows distinct variances across genders and age groups. The resultant WM engagement, though derived from GM, is consistent with metrics measured directly within WM, highlighting a further dependence between GM functionality and WM architecture. Furthermore, the differences observed among groups suggest that WM engagement may serve as a potential biomarker of neurological and cognitive variations, reflecting individual patterns of brain activity and connectivity.

## Methods

### Dataset

Two cohorts from publicly accessible databases were examined: the Human Connectome Project Young Adult (HCP-Y) ^27^ and the Human Connectome Project in Aging (HCP-A)) ^28^. We have included only the relevant MRI modalities that were pertinent to our study from these databases.

### HCP-Y

We selected 687 entries from the HCP-Y repository (comprising 335 males and 352 females, all between the ages of 22 and 35 years), adhering to criteria that included the completeness of a 3T scan and associated physiological data, along with an absence of QC issues. The imaging protocols are detailed elsewhere ^29^ and were performed with 3T Siemens Skyra scanners. Resting-state scans were acquired using multiband gradient-echo EPI sequences. Each session consisted of two runs of scans with opposing phase encoding directions and lasted 14 minutes and 33 seconds, with parameters TR = 720 ms, TE = 33.1 ms, and an isotropic voxel resolution of 2 mm, totaling 1200 volumes. Concurrent recordings of physiological responses, such as respiration and heartbeat, were captured during fMRI scans. Additionally, T1-weighted images were obtained using a 3D MPRAGE sequence with a TR of 2,400 ms, TE of 2.14 ms, and voxel dimensions of 0.7 mm isotropic. T2-weighted images were obtained using a 3D T2-SPACE sequence with a TR of 3200 ms, TE of 565 ms, and voxel dimensions of 0.7 mm isotropic. The diffusion MRI included in our analysis comprised six sequences, each lasting 9 minutes and 50 seconds, using multiband spin-echo EPI, each paired with opposing phase encoding directions. Diffusion weighting consisted of three shells of b=1,000, 2,000, and 3,000 s/mm^2^ interspersed with an approximately equal number of acquisitions on each shell within each run.

### HCP-A

From the HCP-A 2.0 Release, a selection of healthy individuals representing different typical aging cohorts, was analysed ^28,30^. The definition of typical pertains to those showing normal health conditions for their age without known pathological cognitive impairments. A total of 688 participants with intact fMRI datasets and physiological measurements were downloaded, including 303 males and 385 females aged from 36 to 100 years. In our analysis, two sub-groups were further selected: the middle-aged group (305 subjects, age: 40 - 59 y, 125 male and 180 female) and an older group (279 subjects, 60 - 85 y, 134 male and145 female).

The sex difference was not significant between the two groups (Chi-Square test, p = 0.087). For this study, the focus was narrowed to analyzing only the resting-state fMRI data. The imaging protocols are more thoroughly described elsewhere ^30^. Briefly, scans were executed on Siemens 3T Prisma scanners with 32-channel head coils. The resting-state fMRI protocol involved two runs with opposing phase encoding directions, each 6 minutes and 41 seconds, with TR = 800 ms, TE = 37 ms, voxel dimension = 2 mm isotropic, and a total of 488 volumes for each run, while physiological parameters were also documented.

### Preprocessing

The datasets acquired through ICA-FIX, which have regressed out white matter signals, were not used in this study ^31^. Our approach was to employ ‘uncleaned’ images that underwent only the Minimal Preprocessing Pipelines ^32^. Briefly, T1-weighted images were nonlinearly coregistered to MNI space using FNIRT ^33^, with subsequent processing via the Freesurfer suite, resulting in volumetric and cortical surface partitions along with morphometric data ^34^.

Meanwhile, the T2-weighted images were aligned with the native T1-weighted image using 6 degrees of freedom (DOF) and subsequently registered to MNI space along with the T1-weighted images. The fMRI processing encompassed the removal of head movement artifacts, correction of distortions from susceptibility effects using FSL, based on the two runs of data with opposite-phase encoding directions, and then nonlinear registration to MNI space. Further processing steps included regressing out confounding variables including 12 head movement parameters and physiological noise, modeled by the RETROICOR technique ^35^. This preceded the application of linear trend corrections and temporal filtering using a band-pass filter covering the frequency range of 0.01–0.1 Hz. The fMRI data were finally spatially smoothed using a Gaussian kernel with a 4-mm FWHM.

### Reconstruction of WM engagement

The process of reconstructing engagement is graphically depicted in Figure 1a. Initially, brain images are segmented into 90 distinct cortical regions using the Automated Anatomical Labeling (AAL) atlas ^36^. Subsequently, for each voxel within the brain, an SC template is generated employing a population-averaged diffusion profile ^37^, which is derived from 1065 diffusion datasets of young adults from HCP, the same repository from which our study data are sourced. This choice ensures that the SC template to some extent reflects the structural intricacies of our dataset. The SC template is constructed according to the following steps: by setting each voxel as a seed, fiber tracking is executed via the DSIstudio software ^38,39^, which applies the population-averaged diffusion profile to delineate potential white matter tracts that pass through the seed. Consequently, for a given voxel *x*, we assemble a 90×90 SC matrix. Within this matrix, the element *S_x_(i, j)* quantifies the fibers interlinking ROI *i* and *j* that intersect voxel *x*. In a simultaneous procedure, functional connectivity (FC) analysis using fMRI data from each subject required computing the mean time series for each ROI and constructing a 90×90 correlation or FC matrix. The matrix elements represent pairwise Pearson’s correlation coefficients between ROI signals. In the FC matrix, to enhance the signal-to-noise ratio, only correlation values exceeding 0.2 are retained for further examination. Using the brain connectivity toolbox ^38^, the EBC for each neural linkage is calculated, yielding a 90×90 EBC matrix. Edge betweenness centrality quantifies the frequency with which an edge occurs on all shortest paths within a network. Therefore, the element *E(i, j)* signifies the functional significance of the connection between ROIs *i* and *j* with respect to the entire neural network.

**Figure 1:**
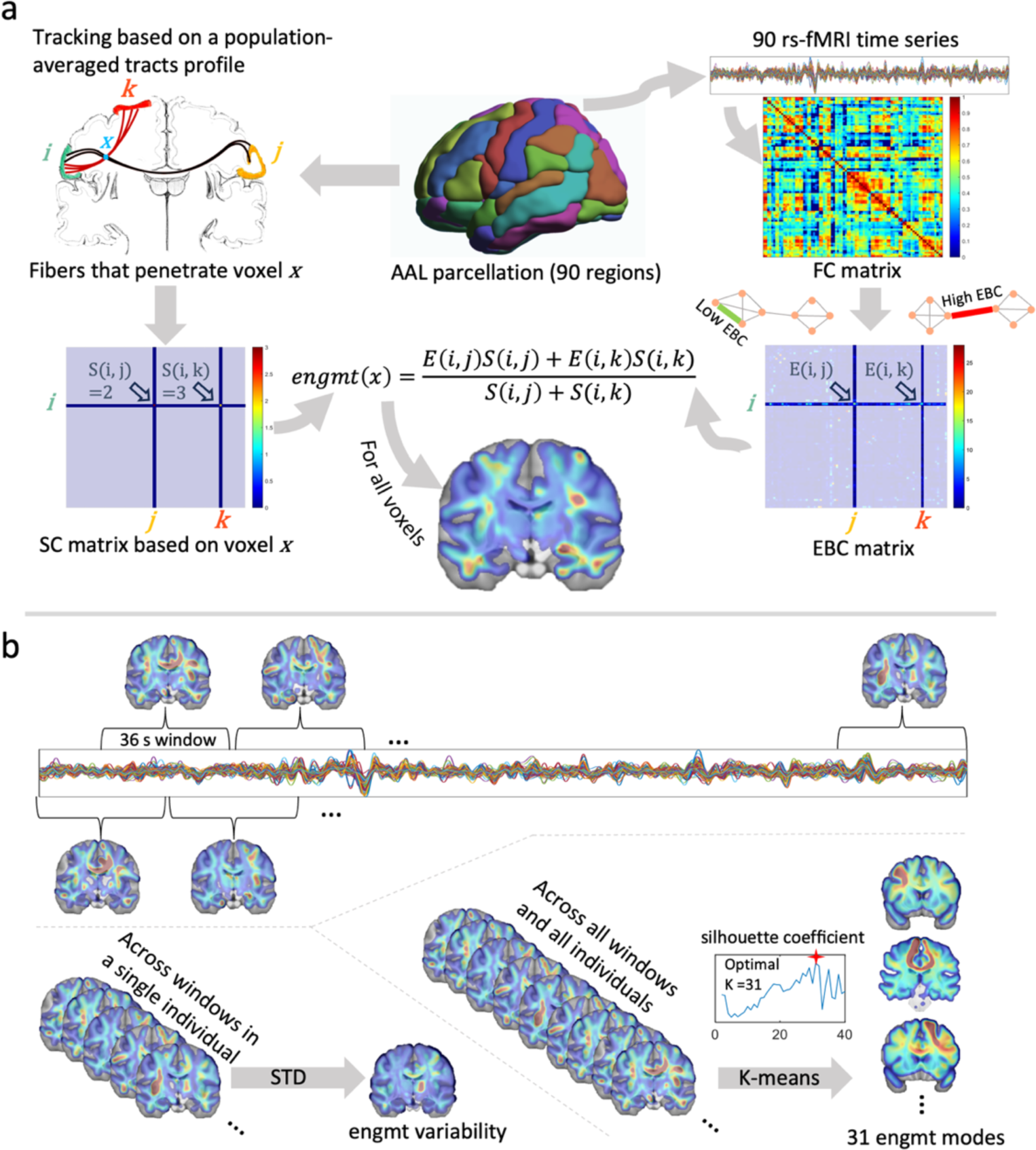
Reconstruction and Analysis of Dynamic White Matter Engagement. **1a**: Workflow of WM engagement reconstruction. The brain is parcellated into 90 GM regions using the AAL atlas, and fiber tracking is performed from each voxel as a seed point to create an SC matrix, indicating the number of fibers connecting pairs of regions through each voxel. FC is calculated from the fMRI time series of the 90 regions, producing an FC matrix based on Pearson’s correlation coefficients. The EBC matrix is derived to represent the functional importance of each connection. Voxel engagement (engmt) is then computed as a weighted average of the EBC matrix, with the SC matrix serving as weights. **1b**: Dynamic Engagement Analysis. To capture the temporal dynamics of voxel engagement, the time course is divided into overlapping windows of 36 seconds. The standard deviation (STD) of voxel engagement across these windows quantifies temporal variability. The clustering of concatenated engagement maps from all windows and subjects via k-means results in 31 mode maps, each representing a cluster center. The silhouette coefficient method determines the optimal number of clusters. Mode occurrence is measured by the frequency of each mode map across the time course for each subject, reflecting the dynamic engagement of voxels in the network.

The final stage involves computing the engagement value for each voxel *x*. This is achieved by taking a weighted average of the EBC matrix elements, where the weights correspond to the equivalent elements in the SC template matrix *S_x_*. Through this method, it becomes evident that a voxel situated on the structural conduits connecting nodes with substantial influence on network communication is deemed to have high engagement. In addition, non-zero *S_x_* could be identified in only high-diffusion areas, so GM voxels usually exhibit low or no engagements.

### Dynamic engagement

Functional neural networks may change dynamically so we hypothesize that voxel engagement is also not static but changes over time, influenced by the network characteristics from which it is derived. As illustrated in Figure 1b, engagement metrics may be computed over sliding time windows instead of over the entire time course. We defined time windows of length 36 seconds with 18-second overlaps, which aligns with established recommendations ^39,40^. Using sliding windows we evaluated the temporal variations of engagement, and the standard deviation of engagement values across all windows was used as a quantitative measure of this variability. To further delve into the dynamics, we concatenated engagement maps derived from each window and across all participants, and subjected them to k-means clustering. This process organizes the maps into 31 distinct clusters, with the centroid of each cluster defined as a mode. The selection of 31 clusters is informed by the silhouette coefficient, a measure of cluster validity, maximized over a range of cluster counts (k = 1 to 40). Subsequently, we quantify the occurrence of each mode within the data of each individual by counting how frequently each mode appears throughout the time course.

### Along-tract analysis of the engagement map

The along-tract analysis enables a detailed exploration of how engagement varies along specific, well-defined white matter tracts, potentially reducing the data dimensions of the original engagement maps while also improving the signal-to-noise ratio by averaging over voxels with similar structural characteristics. We employ a population-averaged tractography atlas ^37^, which includes 70 major WM tracts constructed from HCP young adult data. For each tract containing *m* fibers of varying lengths, we evenly sample 100 points per fiber. The engagement values at these points are compiled into an *m×100* matrix. This matrix was then averaged across *m* rows to create a representative engagement profile for the tract. These vectors enable tract-wise engagement comparisons by performing t-tests individually on the 100 elements between groups. Statistical rigor is maintained by applying False Discovery Rate (FDR) correction methods to account for the multiple comparisons inherent in this analysis. Moreover, the tract atlas also categorizes tracts into associative, projection, and commissural groups. This categorization allows for nuanced comparisons of engagement across diverse WM architectures.

### Structural and functional measurements in WM

To explore the possible structural, functional, and biophysical implications of WM engagement, we analyze its relationship to a suite of imaging parameters that are measured directly from WM. These parameters include myelin content, fiber orientation dispersion (OD), regional homogeneity (Reho), fractional amplitude of low-frequency fluctuations (fALFF), and non-neurite cell density (1-NDI).

### Myelin content

We quantified myelin content using the T1-weighted (T1w) to T2-weighted (T2w) ratio method, leveraging data obtained from the HCP young adults. This approach is grounded in the principle that the ratio of T1w to T2w signal intensities in brain images correlates with myelin content, as demonstrated by Glasser and Van Essen ^41^ and further elaborated by Ganzetti et al. ^42^. The T1w and T2w images were registered into MNI space so that inter-scan movement and anatomical differences were removed. Following this, the T1w/T2w ratio was computed for each voxel, resulting in a map used to depict myelin distribution and density across different brain regions.

### Reho

ReHo is a measure that evaluates the similarity of neuronal activity patterns within a cluster of neighboring voxels ^43^. Kendall’s coefficient of concordance (KCC) is used to assess this similarity, with values ranging from 0 to 1, where higher values indicate greater similarity. ReHo does not require predefined ROIs and can provide information about local brain activity. In this study, a cluster size of 27 voxels is used, which means that the calculation includes the given voxel and its immediate 26 neighbors.

### fALFF

fALFF is a measure used to quantify the relative amplitude of low-frequency oscillations (LFOs) in resting-state fMRI data ^44^. It is defined as the power within the low-frequency range of 0.01-0.1 Hz divided by the total power in the entire detectable frequency range, thus representing the proportion of LFOs within the overall frequency spectrum observed in the BOLD signal. The calculation of fALFF involves transforming voxel time series data into the frequency domain using Fourier transformations to compute a power spectrum. The fALFF is then calculated as the sum of the spectral amplitudes within the 0.01-0.1 Hz frequency range divided by the total amplitude across the entire frequency range measured.

### OD and 1-NDI

OD is a parameter derived from diffusion MRI models that quantify the spread of fiber orientations within a voxel. A higher OD value indicates a larger spread of axon orientations, representing more heterogeneous fiber arrangements within the voxel. This measure is derived from the Neurite Orientation Dispersion and Density Imaging (NODDI) model, which provides a set of parametric maps including isotropic volume fraction (ISOVF), intraneurite volume fraction (NDI), and OD. NODDI fitting on each voxel was performed using the Accelerated Microstructure Imaging via Convex Optimization (AMICO) ^45^. NDI is a parameter that quantifies the volume fraction of neurites (such as axons and dendrites) within a voxel. The quantity (1 - NDI) represents the inverse, which is the volume within a voxel not occupied by neuronal structures and thus acts as a surrogate for the remaining volume fraction which in WM is largely comprised of glial cells.

## Results

### Spatial distribution of the engagement map

Figure 2 shows the spatial distribution of engagement across the brain, presented both voxel-wise (2a) and tract-wise (2b). Voxel-wise analysis reveals a pattern of distribution that exhibits a degree of symmetry, with notable peaks of engagement identified within the frontal, temporal, and parietal WM regions as well as the cingulum. The tract-wise distribution further clarifies these findings, with the left cingulum-frontal-parahippocampal tract (C_FPH_L) and the left cingulum-frontal-parietal tract (C_FP) demonstrating the most pronounced engagement across the cerebral terrain. Additionally, the right parietal aslant tract (PAT_R) is distinguished by its uniformly high engagement along its entire length. Moreover, the left superior longitudinal fasciculus 1 (SLF1), encompassing the dorsal segment of frontal-parietal connections, together with the left cingulum parolfactory tract (C_PO_L) and the right arcuate fasciculus (AF_R), also exhibit high engagement levels. A comparative analysis across different tract categories reveals that the association tracts are markedly more engaged than either the projection or commissural tracts. Note that the full name of the tracts can be found in Table S1.

**Figure 2.**
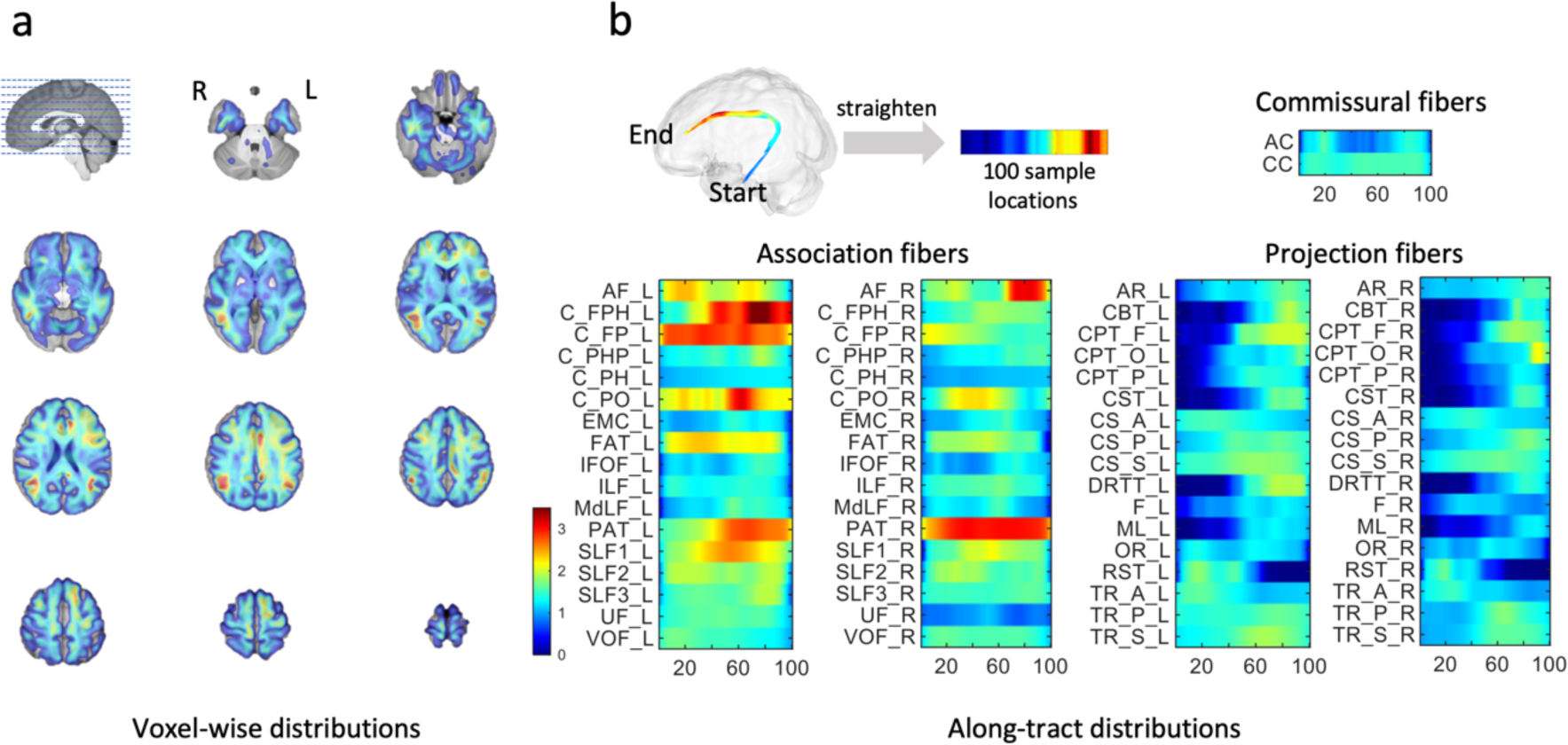
The spatial distributions of measured engagement. **2a**: The voxel-wise distribution of engagement. **2b**: The tract-wise distribution of engagement, categorized into association, projection, and commissural fibers. The full name of the tracts can be found in the supplement information.

### Relationship between engagement and imaging parameters directly measured from WM

Engagement maps for each subject were converted to z-score maps by normalizing with the mean and standard deviation. Subsequently, a one-sample t-test identified clusters exhibiting significantly high and low engagement at the group level, as compared to the global mean activity of zero on the z-map. Figure 3a displays these clusters, indicating regions of both significant high and low engagement throughout the brain (p<0.001, FWE correction). These regions of interest were then used to extract average imaging parameters that have been measured directly from WM, and pairwise t-tests compared these parameters between the two ROIs across all subjects. Figures 3b-f present violin plots for myelin content, ReHo, OD, fALFF, and 1-NDI within the ROIs. In the high-engagement areas, a significant increase in myelin content was observed, suggesting higher efficiency in neural signal conduction. Elevated ReHo in these regions implies more robust local FCs. In contrast, OD values were significantly lower in high-engagement areas, indicating a potential decrease in fiber orientation dispersion. Additionally, the high-engagement regions exhibited significantly greater fALFF, reflecting the intensity of intrinsic neural activities. Lastly, higher 1-NDI values in these areas suggest a denser population of non-neurites, including glial elements, in the high-engagement regions.

**Figure 3.**
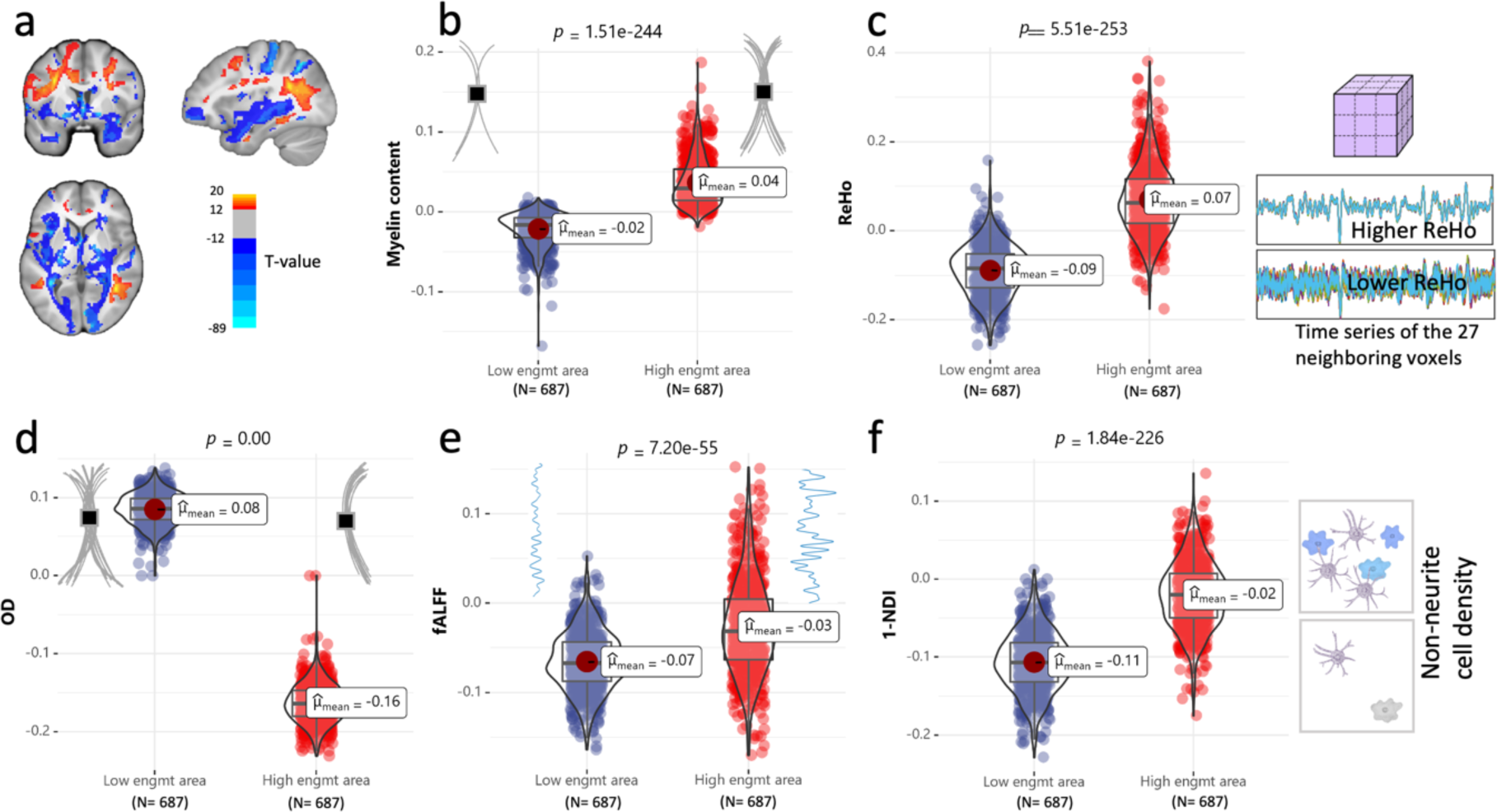
Relationship between engagement and imaging parameters directly measured from WM. **3a**: Spatial distribution voxels that show significant high and low engagement (p<0.001, FWE correction). The red and blue areas respectively indicate the high and low –engagement areas from which the imaging parameters in 3b-f are measured. **3b-f**: Comparison of myelin content, Reho, OD, fALFF, and 1-NDI between the high-engagement areas (red) and low-engagement areas (blue) across all subjects. The statistical result from the pair-wise t-test is indicated above each violin graph.

### Reproducibility of the engagement map

An assessment of reproducibility was conducted through a bifurcation of the subject pool into two distinct groups at random, followed by a comparative analysis of their corresponding engagement maps. As delineated in Figure 4, the visualization offers a detailed depiction of along-tract engagement on a tract basis. The sections of tracts that are anticipated to demonstrate significant differences between the groups were intended to be highlighted by shading. However, the absence of such shading across the panels signifies a lack of statistically significant differences in engagement between the two randomly formed groups. This uniformity is indicative of high reproducibility across the engagement maps, suggesting that the engagement measure is stable and not subject to substantial variance between different subsets of the population.

**Figure 4.**
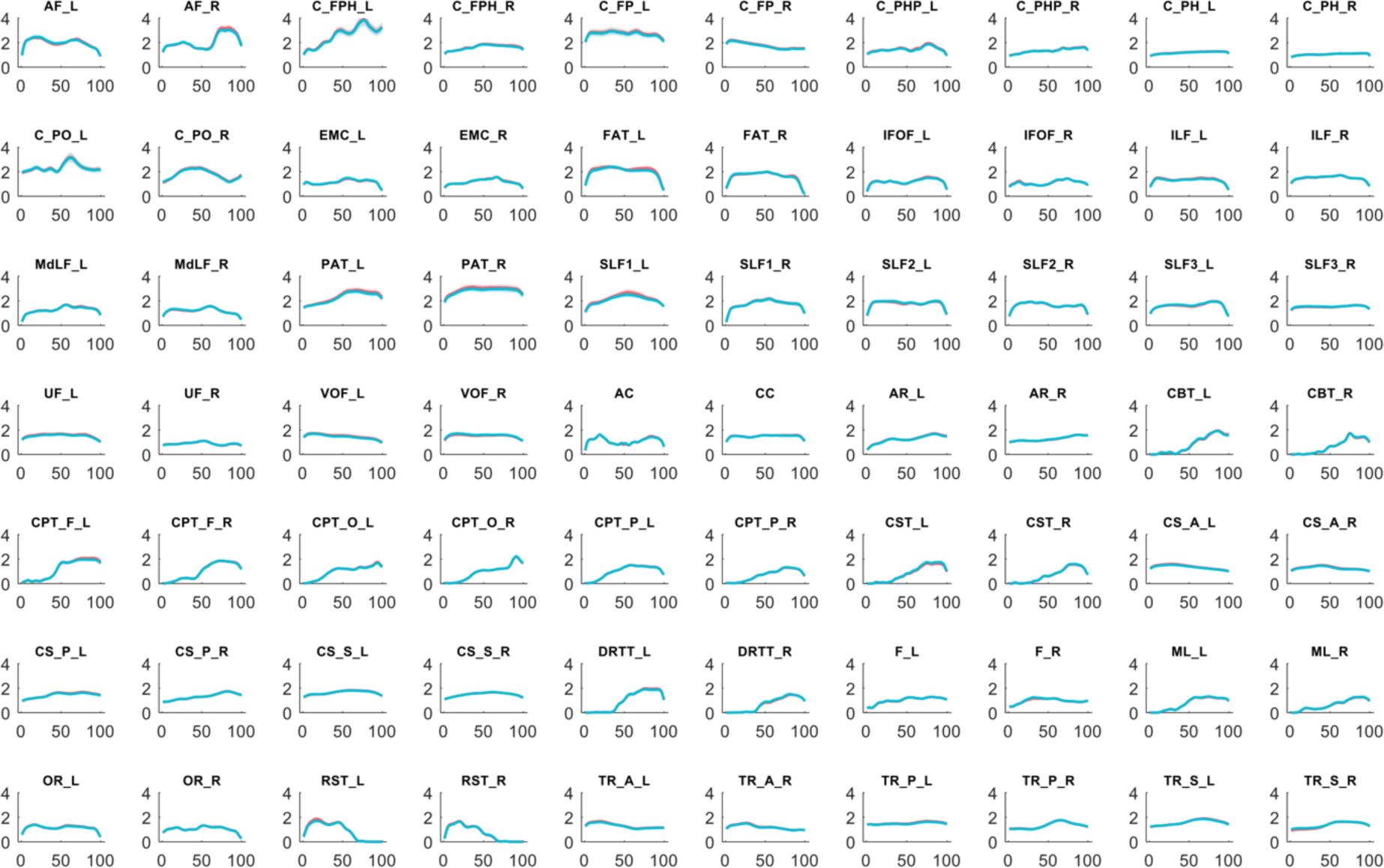
Comparison of along-tract engagement between two random groups. Each panel represents a tract, with the mean along-tract engagement for the two groups delineated by solid lines in blue and red. The shaded bands around these lines denote the 95% confidence intervals, encapsulating the expected range of variation in engagement values if the experiment were repeated. The x-axis of each panel maps the sampled points along the tract, while the y-axis quantifies the degree of engagement. Notably, the sections of tracts that are anticipated to demonstrate significant differences between the groups were intended to be highlighted by shading (p<0.05, FDR correction). The uniformity across panels, with no significant shading differences, indicates a reproducible engagement measure across both cohorts.

### Comparison of engagement between different sexes

Figure 5 shows the mean engagement for each gender, with males represented by blue lines and females by red lines. From the figure, 56 out of 70 tracts exhibit significant differences (p<0.05, FDR correction) in engagement between females and males. Tracts including C_FPH_R, C_FP_L, C_PH_L, FAT_L, SLF1_L, CC, CS_S_L, CS_S_R, TR_P_L, and TR_S_L show a consistent separation of mean engagement levels between genders across the entire sampled length, suggesting a robust gender-specific difference in engagement within these tracts. Conversely, tracts such as C_PH_R, PAT_L, UF_L, UF_R, VOF_L, VOF_R, AC, CBT_R, CS_A_R, F_R, ML_R, OR_R, RST_R, and TR_A_R appear to have no gray shading, indicating no significant difference in engagement between genders in these tracts. A general observation is that in tracts with significant differences, females consistently exhibit higher engagement levels than males. No tracts were found where male engagement surpassed that of females.

**Figure 5.**
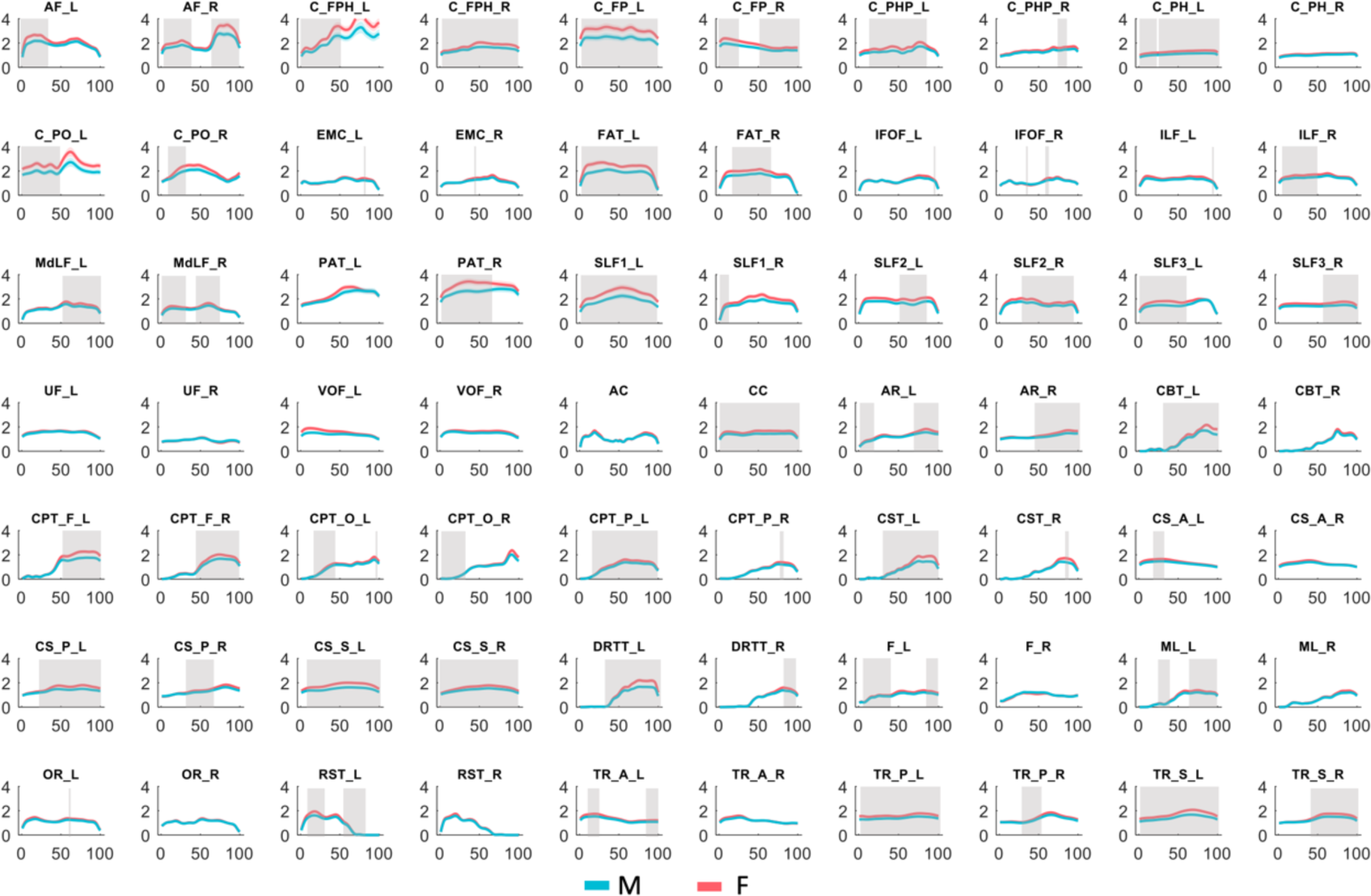
Comparison of along-tract engagement between male and female. Each panel represents a tract, with the mean along-tract engagement for the male and female groups delineated by solid lines in blue and red respectively. The shaded bands around these lines denote the 95% confidence intervals, encapsulating the expected range of variation in engagement values if the experiment were repeated. The x-axis of each panel maps the sampled points along the tract, while the y-axis quantifies the degree of engagement. Notably, the sections of tracts that show significant differences between the groups are highlighted by gray shading (p<0.05, FDR correction).

### Comparison of engagement between different age groups

Figure 6 compares along-tract neural engagements between middle-aged and elderly groups from the HCP-A data. Notably, discernible variances are confined to segmented regions within the AF_L and the EMC_R, where the disparities reach statistical significance (p<0.05, following FDR correction). This finding suggests a higher vulnerability to age-related changes in these tract segments. Although some tracts exhibit elevated engagement levels in the middle-aged group compared to the elderly, these differences do not meet the threshold for statistical significance. Overall, while engagement levels appear relatively consistent across the majority of tracts, a trend of higher engagement in the younger cohort can be discerned, suggesting subtle yet pervasive effects of aging on neural connectivity.

**Figure 6.**
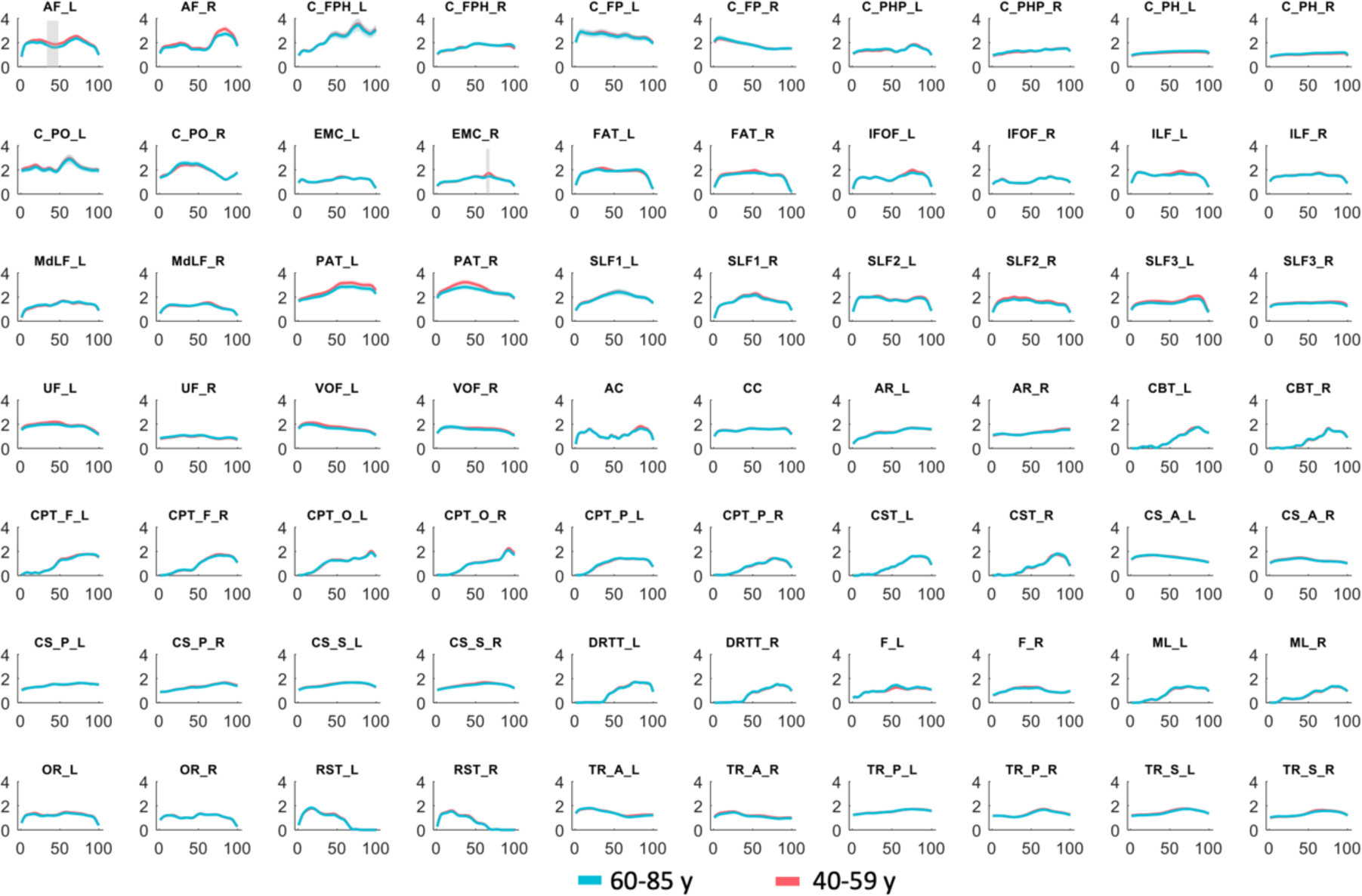
Comparison of along-tract engagement between middle-aged and old groups. Each panel represents a tract, with the mean along-tract engagement for the mid-aged and old groups delineated by solid lines in red and blue respectively. The shaded bands around these lines denote the 95% confidence intervals, encapsulating the expected range of variation in engagement values if the experiment were repeated. The x-axis of each panel maps the sampled points along the tract, while the y-axis quantifies the degree of engagement. Notably, the sections of tracts that show significant differences between the groups are highlighted by gray shading (p<0.05, FDR correction).

### The distribution and occurrences of the engagement modes

Figure 7a depicts 31 distinct neural engagement patterns, identified through a clustering analysis of window-wise engagement maps, with modes 4 and 24 shaded to denote their exclusion due to identified spatial artifacts. It is noteworthy that the majority of these modes correspond to the anatomical distribution of white matter tracts. For instance, mode 3 distinctly mirrors a segment of tracts traversing the corpus callosum, facilitating interhemispheric connectivity, while mode 30 is reminiscent of the anterior corona radiata. Panel 7b presents a quantitative comparison of mode occurrences between sexes, highlighted by blue bars for males and red for females. Our analysis reveals that modes 8, 13, 14, 15, and 29 are more prevalent in females, displaying symmetry (8 vs. 13, and 14 vs. 15) and predominantly representing connections involving bilateral motor areas, the bilateral frontal aslant tract, and the cingulum. Intriguingly, mode 17 registers as the most frequent across sexes, with a higher incidence in males. This mode’s distribution is unique, characterized by a consistently low engagement throughout the brain.

**Figure 7.**
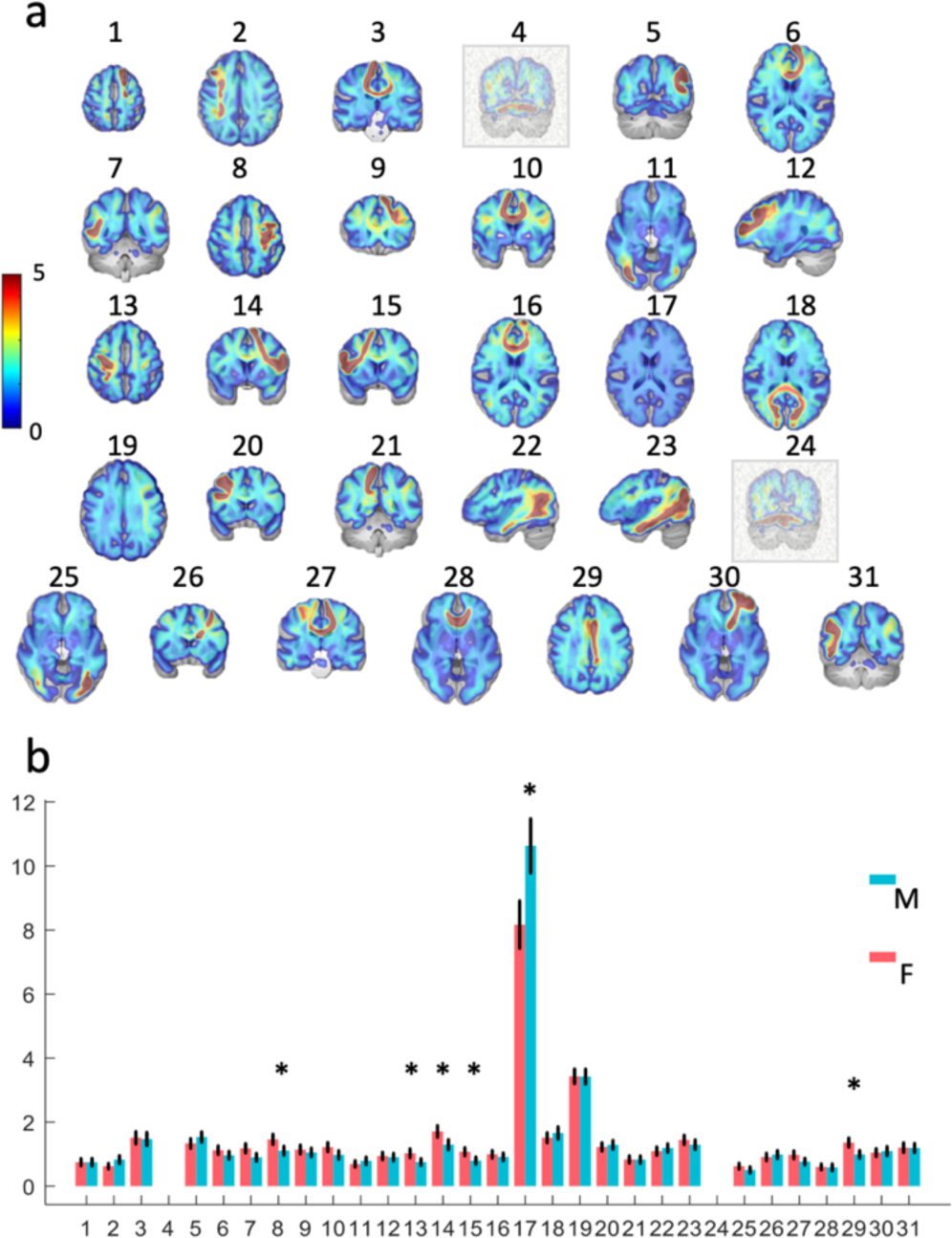
Spatial distribution and occurrences of the engagement modes. **7a**: Thirty-one engagement modes mapped across the brain, identified through clustering of engagement maps across time windows and individuals. Modes 4 and 24 are shaded and excluded due to artifacts. **7b**: Occurrences of engagement modes in males (blue) and females (red), with the bar height representing the group mean and error bars indicating the 95% CI. An asterisk (*) denotes significant differences between sexes (p<0.05, Bonferroni corrected). Modes 8, 13, 14, 15, and 29 show higher female occurrences, while mode 17 is more common in males and exhibits a unique low engagement pattern.

### Engagement variability over time

Figure 8a illustrates the spatial distribution of engagement variability across time, averaged across participants. This map, distinct in its characteristics from the engagement map, shows bilateral symmetry and pronounced variabilities proximal to gray matter regions and along tracts traversing the corpus callosum. In Figure 8b, we observe gender-specific patterns of mean engagement variability, delineated by blue lines for males and red lines for females. A notable gender disparity is evident in 58 of the 70 examined tracts, where significant differences (p<0.05, FDR corrected) in engagement variability were found. Tracts including C_FP_L, FAT_L, SLF1_L, CC, and CS_S_L show consistent differences between genders across the entire sampled length, suggesting a robust gender-specific difference in engagement variability within these tracts. In contrast, tracts including C_PHP_L, C_PHP_R, C_PH_L, C_PH_R, PAT_L, SLF2_R, VOF_L, VOF_R, CS_A_R, F_L, OR_R, and TR_A_R showed no significant differences in engagement variability between the genders. The overarching trend indicates that females tend to exhibit higher variability in engagement compared to males.

**Figure 8.**
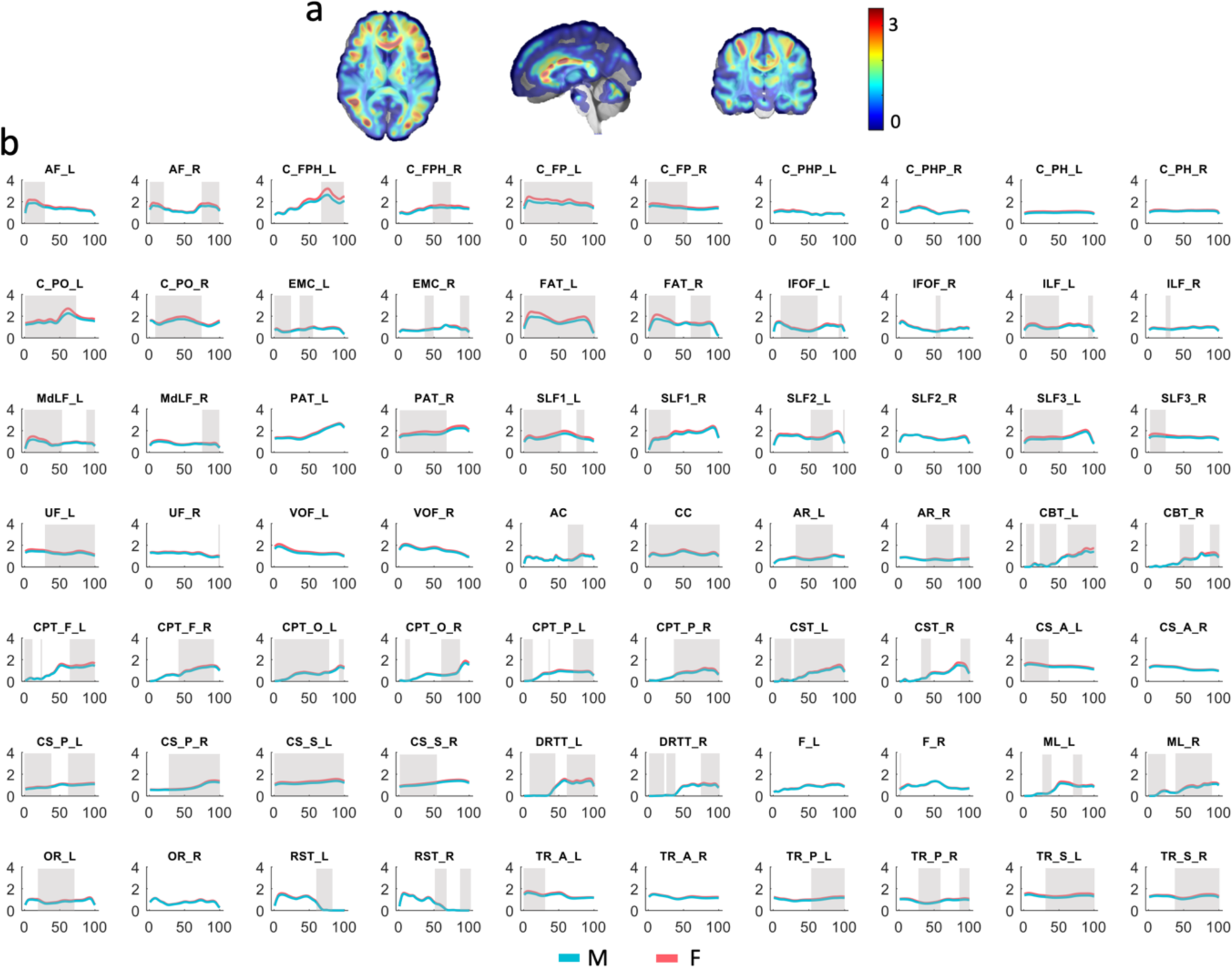
Visualization of neural engagement variability and gender-specific differences. **8a**: The spatial distribution of neural engagement variability, averaged across individuals and visualized in a brain map. This map highlights areas of high variability, particularly near gray matter and across the corpus callosum. **8b**: Comparison of along-tract engagement variability between males and females. Each panel represents a tract, with the mean along-tract engagement variability for the male and female groups delineated by solid lines in red and blue respectively. The shaded bands around these lines denote the 95% confidence intervals. The x-axis of each panel maps the sampled points along the tract, while the y-axis quantifies the degree of engagement variability. Notably, the sections of tracts that show significant differences between the groups are highlighted by gray shading (p<0.05, FDR correction).

## Discussion

Our study introduces a novel functional measure of WM engagement, which maps the functional importance of an edge to its structural counterparts. This mapping yields a spatial distribution of engagement that is not only highly reproducible but also aligns with direct structural, functional, and energetic measures within WM, illustrating a distinct inter-dependence between the functions of GM and the characteristics of WM. Additionally, our analysis has uncovered 31 unique neural engagement patterns through a clustering analysis of window-wise engagement maps, highlighting the temporal variability of neural engagement. Our results also uncover significant differences in engagement levels and their variabilities across genders and age cohorts, suggesting that WM engagement, along with its variability identified in our study, could serve as a biomarker of neurological and cognitive diversity, potentially mirroring the unique patterns of brain activity and connectivity in individuals.

The engagement map yields novel insights into the degree of WM involvement in sustaining functional brain networks. The symmetrical pattern observed in the voxel-wise engagement distribution suggests a balanced participation of these regions in neural processing and connectivity. The highest engagement identified in various cingulum-associated tracts corroborates with previous neuroanatomical research underscoring the significance of the cingulum in connecting key areas like the prefrontal cortex and posterior cingulate cortex, both crucial components of the resting-state default network which is highly active during a resting state ^46^. Further, the involvement of the cingulum in cognitive, affective, and memory-associated functions is well-recognized ^47,48^, reinforcing its vital role delineated by our findings. Importantly, our observations indicate that areas with substantial engagement are predominantly associated with frontal-parietal connections. This includes the previously discussed cingulum, the parietal aslant tract that links the inferior frontal gyrus (encompassing Broca’s area integral to language production) with the posterior parietal lobe (related to motor planning), and the superior longitudinal fasciculus 1, which spans the dorsal segment of the frontal-parietal axis, all demonstrating high engagement levels. These results collectively affirm the critical role of frontal-parietal connections in supporting brain functions. In addition, the observed trend of association tracts being more engaged than projection tracts resonates with the intrinsic nature of our methodology. In our analytical framework, we consider only those voxels that form interconnecting links between pairs of GM regions as edges. Thus, the engagement, also conceptualized as a weighted average of EBC, is more likely to be measurable from association and commissural tracts that connect GM regions than a subset of the projection tracts, such as the cortico-spinal tract, that connect GM and brainstem. This observation may also reflect that association tracts are essential for higher-order cognitive functions, whereas projection tracts, which convey information between the cortex and sub-cortical regions or body parts, are required for lower-order functions, such as visual and motor activities, which are typically less engaged during the resting state when there is no specific sensory input to process or motor output required ^49,50^. Moreover, we noted that association tracts demonstrate greater engagement compared to commissural tracts despite that they both connect GM regions. This may be attributed to the role of association tracts in integrating and processing complex information within the same hemisphere, rather than merely coordinating inter-hemispheric communication. This delineation of engagement patterns holds potential for future research into the functional connectivity of the brain, offering a promising avenue for associating these engagement levels with diverse cognitive processes and neurological conditions.

The comparison of imaging parameters between high and low-engagement areas provides several novel insights. Firstly, the observed increment in myelin content within high-engagement regions suggests they have enhanced axonal conductivity and, by extension, more efficient neural signal transmission ^51^. This aligns with the myelination hypothesis, which posits that high cognitive demands may stimulate oligodendrocyte proliferation and myelin reinforcement ^52^.

Secondly, elevated ReHo in these areas supports the concept that synchronized local neural activity is characteristic of brain engagement, as exemplified by the observation of higher ReHo in the motor cortex during a finger motion task compared to a resting state ^43^. Interestingly, our analysis indicated a significant decrease in OD values in high-engagement areas, suggesting a potential simplification in the complexity or diversity of fiber orientations. This might reflect a more streamlined architectural design, optimized for functioning as a bottleneck conduit in neural networks. This observation dovetails with the ReHo findings, where higher ReHo is indicative of a well-organized, functionally uniform neighborhood of neural activity. The significant augmentation of fALFF in high-engagement regions further suggests that the intensity of intrinsic neural activities is greater in these areas. This enhancement in fALFF could be indicative of increased neural ‘preparedness’ or readiness in response to cognitive demands. Lastly, higher 1-NDI values in high-engagement areas might reflect a denser population of non-neurites, predominantly glial elements, which are increasingly recognized for their role in cognitive functions ^53^. This aligns with our prior observations linking higher 1-NDI in white matter with increased metabolic demand, as evidenced by features of the BOLD hemodynamic response function (HRF) ^54^. The implications of these findings are multifaceted, suggesting a robust structure-function association, as indicated by the relevance between the engagement mapped from GM function and specific neurobiological indices measured in WM. This correlation holds potential for clinical applications, particularly in identifying functional anomalies that are not directly detectable in WM images. However, it is crucial to note the limitations of this study, specifically its reliance on indirect measures of neural components like myelin content and non-neurites. These should be regarded as proxies for the underlying microstructure, necessitating a cautious approach to interpretation. Future research should aim to validate these findings through animal model studies or the use of additional MRI methods.

The findings of this study highlight significant sex differences in neural engagement across most white matter tracts, notably within the cingulum and frontal aslant tract. These disparities are in line with existing research that underscores sex-specific patterns in brain connectivity, such as enhanced connectivity within the Default Mode Network (DMN) observed in females ^55,56^, with cingulum playing a pivotal role in linking key DMN components. The higher engagement in the frontal aslant tract is consistent with our finding regarding higher occurrences of modes 14 and 15 in females than males. These two modes highly resemble the distribution of the frontal aslant tract. Additionally, the age-related differences we observed in the Left Arcuate Fasciculus and Right and Extreme Capsule accord with literature pointing to the selective impact of aging on specific brain regions. The role of Arcuate Fasciculus in language and cognitive functions is well-documented ^57^, and its degradation with age could correspond to the observed decline in language abilities. Despite this, we noticed a relative consistency in engagement levels between two age groups across most tracts, with only a slight increase in middle-aged individuals in small sections of specific tracts, suggesting a potential insensitivity of the engagement to aging. This could be attributed to brain plasticity and compensatory mechanisms within the aging brain^58^. Our data particularly emphasize the overarching influence of sex on neural engagement compared to age, indicating that sex differences in brain structure and function are more pronounced and ingrained than age-related changes. Understanding these sex-specific neural engagement patterns may be crucial as they hold significant implications for sex-specific vulnerabilities to neurological and psychiatric disorders.

Our study elucidates the dynamic nature of white matter (WM) engagement within the brain, revealing a complex pattern that is far from static or random. Through the decomposition of the engagement map into individual modes, we observed a reflection of the architectures of WM tracts. Distinct groups of WM voxels along specific tracts show synchronized engagement for certain periods, indicative of a structured and temporal coordination in their engagement. This dynamic is akin to cortical co-activation patterns (CAPs) ^59^, yet it distinctively illustrates the supportive interactions among WM voxels within the vast neural network. Further probing into the temporal aspects of these engagement modes has unveiled a predominant mode characterized by a globally low level of brain engagement. This prevalent mode, which we interpret as the ‘default baseline state’, suggests a foundational level of activity where the brain may remain for protracted periods. Notably, our findings indicate a gender-specific pattern: males generally sustain this baseline state for a longer duration than females, who, conversely, show more instances of engagement in modes associated with specific WM tracts. Such group-related differences in dynamic WM engagement may hold significant potential as biomarkers for identifying distinct neurological conditions or diseases.

In conclusion, this report introduces an innovative functional measure of WM engagement, mapping the functional significance of neural connections to their structural correlates. This mapping produces a spatial engagement distribution that is not only reproducible but also consistent with structural, functional, and energetic profiles within WM, highlighting a remarkable interdependence between gray matter function and white matter attributes. Differences in these engagement patterns across different genders and ages hint at their potential as biomarkers for neurological and cognitive disorders. This paves the way for future studies to deepen our comprehension of WM engagement, providing insight into brain functionality and contributing to the diagnosis and management of neurological conditions.

## Acknowledgments

This work was supported by the National Institutes of Health (NIH) grants R01 NS113832 (JG), R01 MH123201 (JG), R01 NS129855 (ZD), and K01 EB032898 (KS).

Data of young adults were provided by the Human Connectome Project, WU-Minn Consortium (Principal Investigators: David Van Essen and Kamil Ugurbil; 1U54MH091657) funded by the 16 NIH Institutes and Centers that support the NIH Blueprint for Neuroscience Research; and by the McDonnell Center for Systems Neuroscience at Washington University.

Data of the aging population reported in this publication was supported by the National Institute On Aging of the National Institutes of Health under Award Number U01AG052564 and by funds provided by the McDonnell Center for Systems Neuroscience at Washington University in St.

Louis. The HCP-Aging 2.0 Release data used in this report came from DOI: 10.15154/1520707.

---

**Table S1.** Abbreviations of the tract names.

| Association |  | Projection |  |
| --- | --- | --- | --- |
| AF_L | Arcuate_Fasciculus_L | AR_L | Acoustic_Radiation_L |
| AF_R | Arcuate_Fasciculus_R | AR_R | Acoustic_Radiation_R |
| C_FPH_L | Cingulum_Frontal_Parahippocampal_L | CBT_L | Corticobulbar_Tract_L |
| C_FPH_R | Cingulum_Frontal_Parahippocampal_R | CBT_R | Corticobulbar_Tract_R |
| C_FP_L | Cingulum_Frontal_Parietal_L | CPT_F_L | Corticopontine_Tract_Frontal_L |
| C_FP_R | Cingulum_Frontal_Parietal_R | CPT_F_R | Corticopontine_Tract_Frontal_R |
| C_PHP_L | Cingulum_Parahippocampal_Parietal_L | CPT_P_L | Corticopontine_Tract_Parietal_L |
| C_PHP_R | Cingulum_Parahippocampal_Parietal_R | CPT_P_R | Corticopontine_Tract_Parietal_R |
| C_PH_L | Cingulum_Parahippocampal_L | CPT_O_L | Corticopontine_Tract_Occipital_L |
| C_PH_R | Cingulum_Parahippocampal_R | CPT_O_R | Corticopontine_Tract_Occipital_R |
| C_PO_L | Cingulum_Parolfactory_L | CS_A_L | Corticostriatal_Tract_Anterior_L |
| C_PO_R | Cingulum_Parolfactory_R | CS_A_R | Corticostriatal_Tract_Anterior_R |
| EMC_L | Extreme_Capsule_L | CS_S_L | Corticostriatal_Tract_Superior_L |
| EMC_R | Extreme_Capsule_R | CS_S_R | Corticostriatal_Tract_Superior_R |
| FAT_L | Frontal_Aslant_Tract_L | CS_P_L | Corticostriatal_Tract_Posterior_L |
| FAT_R | Frontal_Aslant_Tract_R | CS_P_R | Corticostriatal_Tract_Posterior_R |
| IFOF_L | Inferior_Fronto_Occipital_Fasciculus_L | CST_L | Cortico_Spinal_Tract_L |
| IFOF_R | Inferior_Fronto_Occipital_Fasciculus_R | CST_R | Cortico_Spinal_Tract_R |
| ILF_L | Inferior_Longitudinal_Fasciculus_L | DRTT_L | Dentatorubrothalamic_Tract_L |
| ILF_R | Inferior_Longitudinal_Fasciculus_R | DRTT_R | Dentatorubrothalamic_Tract_R |
| MdLF_L | Middle_Longitudinal_Fasciculus_L | F_L | Fornix_L |
| MdLF_R | Middle_Longitudinal_Fasciculus_R | F_R | Fornix_R |
| PAT_L | Parietal_Aslant_Tract_L | ML_L | Medial_Lemniscus_L |
| PAT_R | Parietal_Aslant_Tract_R | ML_R | Medial_Lemniscus_R |
| SLF1_L | Superior_Longitudinal_Fasciculus1_L | OR_L | Optic_Radiation_L |
| SLF1_R | Superior_Longitudinal_Fasciculus1_R | OR_R | Optic_Radiation_R |
| SLF2_L | Superior_Longitudinal_Fasciculus2_L | RST_L | Reticulospinal_Tract_L |
| SLF2_R | Superior_Longitudinal_Fasciculus2_R | RST_R | Reticulospinal_Tract_R |
| SLF3_L | Superior_Longitudinal_Fasciculus3_L | TR_A_L | Thalamic_Radiation_Anterior_L |
| SLF3_R | Superior_Longitudinal_Fasciculus3_R | TR_A_R | Thalamic_Radiation_Anterior_R |
| UF_L | Uncinate_Fasciculus_L | TR_P_L | Thalamic_Radiation_Posterior_L |
| UF_R | Uncinate_Fasciculus_R | TR_P_R | Thalamic_Radiation_Posterior_R |
| VOF_L | Vertical_Occipital_Fasciculus_L | TR_S_L | Thalamic_Radiation_Superior_L |
| VOF_R | Vertical_Occipital_Fasciculus_R | TR_S_R | Thalamic_Radiation_Superior_R |
| Commissural |  |  |  |
| AC | Anterior_Commissure |  |  |
| CC | Corpus_Callosum |  |  |

